# Bimodal occupancy-frequency distributions uncover the importance of regional dynamics in shaping marine microbial biogeography

**DOI:** 10.1101/039883

**Authors:** Markus V. Lindh, Johanna Sjöstedt, Börje Ekstam, Michele Casini, Daniel Lundin, Luisa W. Hugerth, Yue O. O. Hu, Anders F. Andersson, Agneta Andersson, Catherine Legrand, Jarone Pinhassi

**Author notes:** To whom correspondence should be addressed. Centre for Ecology and Evolution in Microbial model Systems - EEMiS, Linnaeus University, Barlastgatan 11, SE-39182 Kalmar, Sweden. Author contributions: M.V.L., C.L. and J.Pi. conceived the study; M.V.L., M.C., C.L. and J.Pi. designed research; M.V.L., J.S., L.H. and Y.H. performed research; M.V.L., J.S. B.E., D.L., L.H., A.F.A., A.A. and J.Pi analyzed data; M.V.L., J.S. B.E. and J.Pi. wrote the paper. All authors discussed the results and commented on the manuscript.

## Abstract

Metapopulation theory developed in terrestrial ecology provides applicable frameworks for interpreting the role of local and regional processes in shaping species distribution patterns. Yet, empirical testing of metapopulation models on microbial communities is essentially lacking. Here we determined regional bacterioplankton dynamics from monthly transect sampling in the Baltic Sea Proper (16 sites, 11 occasions, 2010-2011) using 16S rRNA gene pyrosequencing. A strong positive correlation was found between local relative abundance and occupancy of populations. Notably, the occupancy-frequency distributions (the number of populations occupying different number of sites) were significantly bimodal with a satellite mode of mostly rare endemic populations and a core mode of abundant cosmopolitan populations (e.g. *Synechococcus*, SAR11 and SAR86 clade members). Observed temporal changes in population distributions supported theoretical predictions that stochastic variation in local extinction and colonization rates accounted for observed bimodality. Moreover, bimodality was found for bacterioplankton across the entire Baltic Sea, and was also frequent in globally distributed datasets where average Bray-Curtis distances were significantly different between bimodal and non-bimodal datasets. Still, datasets spanning waters with distinct physicochemical characteristics or environmental gradients, e.g. brackish and marine or surface to deep waters, typically lacked significant bimodal patterns. When such datasets were divided into subsets with coherent environmental conditions, bimodal patterns emerged, highlighting the importance of positive feedbacks between local abundance and occupancy within specific biomes. Thus, metapopulation theory applied to microbial biogeography can provide novel insights into the mechanisms governing shifts in biodiversity resulting from natural or anthropogenically induced changes in the environment.

**Significance statement:** Marine bacteria regulate global cycles of elements essential to life and respond rapidly to environmental change. Yet, the ecological factors that determine distribution and activity patterns of microbial populations across different spatial scales and environmental gradients remain basically unconstrained. Our metapopulation model-based analyses show that dispersal-driven processes contribute to structuring the biogeography of marine microorganisms from small to large geographical areas. Discovery of bimodal distribution patterns pinpointed satellite microbial populations with highly restricted ranges and defined abundant core populations widely distributed in coherence with environmental conditions. Thus, application of metapopulation models on microbial community structure may allow the definition of biogeographic regions critical for interpreting the outcome of future ocean changes.

**Classification:** Biological Sciences, Environmental Sciences

## Introduction

Marine microorganisms regulate ecosystem services essential to life through their influence on fluxes of matter and energy in the ocean (1). Conspicuously, these fluxes are ever dynamic due to the pronounced potential of microbial communities to rapidly respond to environmental change, both through adjustments in species composition and metabolic activity (2-4). Although microbial species or populations have been inferred to be active in a variety of distinct biogeochemical processes (see e.g. 5-8), very little is known about the ecological factors that contribute to determining biogeographical distribution patterns of these key organisms. Recent developments in molecular genetics methodologies, i.e. high-throughput DNA sequencing technologies, now allow for detailed spatial and temporal investigations of complex assemblages of microorganisms (9, 10). Thus, it is now not only possible to address questions regarding existing taxonomic diversity but also what ecological processes that potentially influence the structure of natural marine microbial communities.

Historically, local communities have been regarded as independent ecological units with individual integrity, determined by local interactions among coexisting species and environmental factors (11). However, over the past decades, the profound influence of regional factors, e.g. dispersal, in structuring local species assemblages has been established (12-15). A positive relationship between local abundance and regional distribution of species is a pattern characteristic for a wide range of organisms and ecosystems, including both macroorganisms and highly diverse phytoplankton and bacterioplankton assemblages (16-18). Although this relationship is considered to be one of the most robust patterns in community ecology, significant variation in the level of correlation between local abundance and regional distribution occurs among observed assemblages, suggesting that different ecological processes contribute to the shape of the relationship (19-21). There is now an increased awareness of the importance of regional dynamics also for the structuring of aquatic bacterioplankton communities (14, 15, 21). Regional dynamics are linked to a second pattern - the species rank-abundance distribution that describes the abundance of a species in an area or sample as a function of its abundance rank. Especially at larger spatial scales, species abundances exhibit a log-normal distribution, where a few species make up a high proportion of all extant individuals while a majority of species are rare (11, 18, 22, 23, 24). High throughput analyses of linkages between spatial and temporal patterns of bacterioplankton populations across biomes, through assessment of relative sequence abundances, have provided thorough knowledge of the vast genetic diversity dwelling beyond the ocean surface (25, 26). These studies establish that prokaryotic assemblages share much of the same characteristics of biogeographic distribution patterns as macroorganisms. Such high-throughput data from biogeographical studies may disentangle whether mechanistic concepts of distribution patterns in terrestrial systems are applicable also to assemblages of marine pelagic prokaryotes.

Two major hypotheses for explaining ecological mechanisms shaping species distribution patterns have attracted important attention; Levin’s model and the core-satellite (CSH) hypothesis (11, 16, 27). When applied in terrestrial ecology, these models and their modified versions have provided explanatory power to identify the drivers of distribution patterns among insects, plants and animals; this has had particular significance for developing strategies in conservation biology (28). In essence, metapopulation theory describes the dynamics of populations in a patchy environment, i.e. how local colonization and extinction rates interact with species distribution at a regional scale. The two original models differ in how colonization and extinction rates are calculated to predict occupancy-frequency distributions (the number of sites occupied by different number of species), and have been elaborated over time (29, 30).

Levin’s model predicts a unimodal occupancy-frequency distribution characterized by a skewed pattern where most species in a region occupy a single site and the number of species decreases with increasing number of sites (27). Hanski’s metapopulation model or the CSH hypothesis (16-18) predicts a quadratic function of both colonization and extinction rates and takes into account positive feedback mechanisms between local abundance and occupancy. Extinction rates are therefore predicted to be low for populations with high and low occupancy but high for populations with intermediate occupancy. In contrast to Levin’s model, the CSH predicts bimodal distributions where the decrease in number of species with increasing number of study sites is followed by an increase in species occupying all or most sites (16). Thus, the predicted occupancy-frequency pattern in CSH is characterized by a mode of many endemic satellite populations present at only a few sites, and a second mode of cosmopolitan core populations present at all sites. It has been suggested that such bimodal occupancy-frequency patterns should be the norm in aquatic habitats, not least for free-living easily dispersible prokaryotes (31). However, there has been little empirical testing of these models in aquatic habitats in general and for aquatic microorganisms in particular (but see (32)).

In the present study, we used metapopulation models to analyze local and regional dynamics of bacterioplankton populations (operational taxonomic units [OTUs], defined at 97% 16S rRNA gene sequence identity). We investigated regional effects on monthly sampled local communities in the Baltic Sea Proper by determining: (i) the relationship between local OTU abundance and OTU distribution, i.e. occupancy, (ii) the type of occupancy-frequency distribution pattern, i.e. the number of OTUs occupying different number of sites, and (iii) the relationship between OTU occupancy and colonization and extinction rates. We further identified OTUs showing distinct distributional and colonization/extinction patterns. To validate observed distribution patterns we extended the analyses of occupancy-frequency distributions to complementary datasets spanning the entire Baltic Sea as well as global International Census of Marine Microbes (ICoMM) datasets. We also evaluated observed OTU distributions with beta-diversity patterns and variability in environmental conditions available as metadata in the ICoMM database.

## Results and Discussion

### Bacterioplankton community composition in the Baltic Sea Proper

We investigated bacterioplankton population dynamics in a grid of 16 stations in the Baltic Sea Proper over two years (2010-2011) to elucidate the role of regional dynamics in shaping biogeographical patterns (Fig. S1). Bacterioplankton community composition showed pronounced clustering of samples from the two years according to season (Fig. S2). This recurrent seasonal succession in bacterioplankton agrees with the recognized importance of seasonal shifts in environmental conditions in shaping community composition of marine bacteria (23, 24, 25). There was also a tendency towards differentiation of community composition between coastal and offshore sites (Fig. S2).

Between 122 and 279 OTUs were detected per sampling point, out of which between 10 and 22 OTUs were abundant while a grand majority (typically 100-257 OTUs) were rare, when defining abundant and rare OTUs as having relative abundances >1% and <0.1%, respectively (33). All sampled communities displayed rank-abundance curves following a log-normal distribution (Fig. S3), typical of aquatic microbial communities, and widespread also among macroorganism communities in terrestrial environments (24, 33-36).

The pattern of many rare compared to a few abundant populations for organisms in a variety of habitats may be inherently linked with strong regional dynamics, i.e. a positive relationship between local abundance and regional occupancy. Such regional dynamics have been established for a wide range of organisms in different ecosystems, including bacterioplankton in aquatic environments (19-21). We therefore plotted the number of sites occupied by specific OTUs versus their average local relative abundance (Fig. 1). We used locally weighted regression spline smoothing (LOESS) to indicate the positive relationship between average local relative abundance of OTUs with average occupancy. This plot also showed that a majority of OTUs had low abundance and low occupancy compared to relatively few OTUs with high abundance and high occupancy. This suggests that the mechanisms and regional dynamics typically observed among macroorganisms outside the marine realm also shape bacterioplankton assemblages in surface waters.

**Figure 1.**
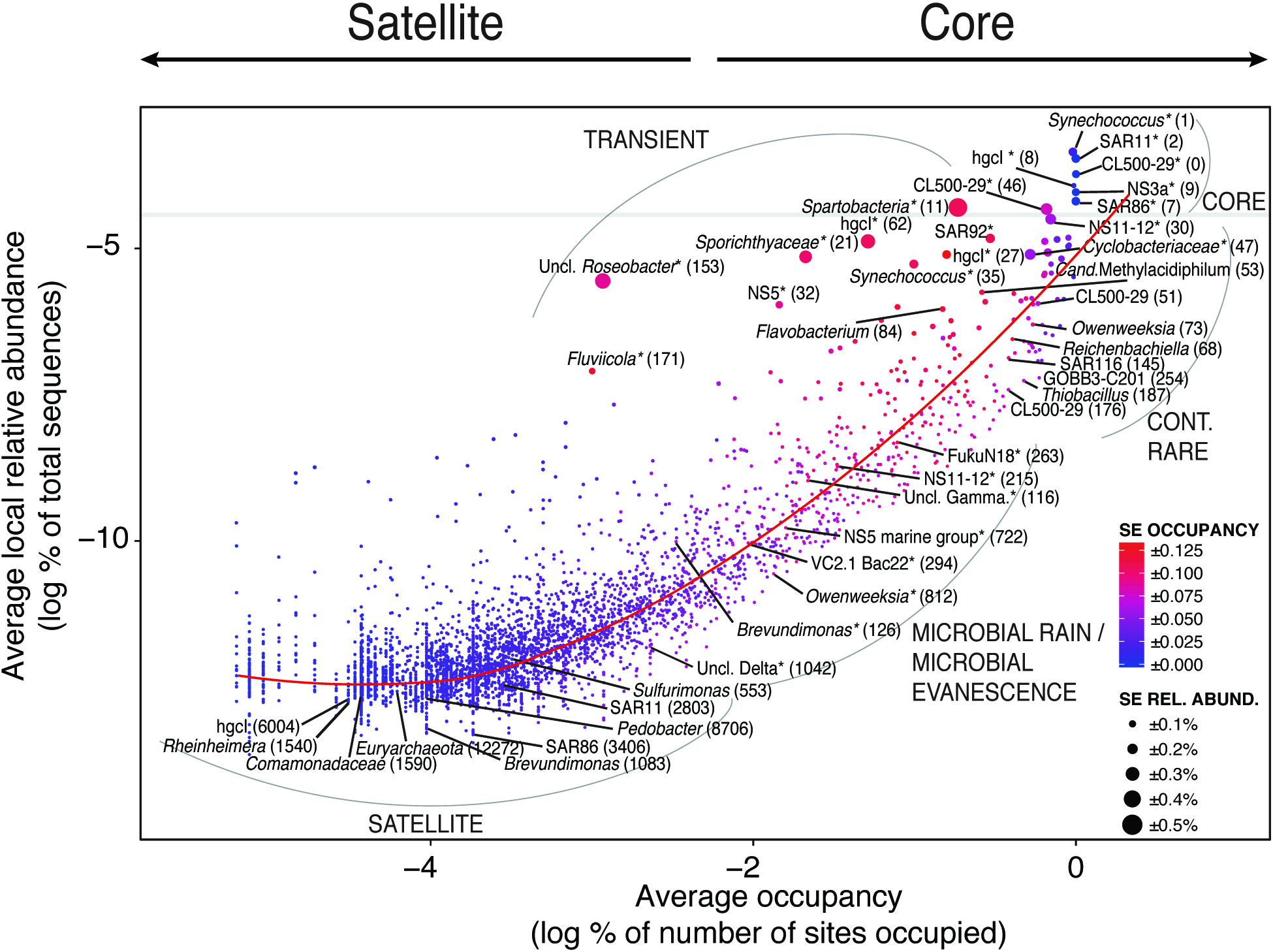
Average relative abundance (normalized sequence abundance) against average occupancy. Color denotes standard error in occupancy where red is high and blue is low variation. Size denotes standard error in relative abundance where large size is high and small size is low variation. By moving towards left on the x-axis OTUs display satellite characteristics and towards right OTUs display core characteristics. Examples of OTUs characterized as core, satellite, transient, continuously rare and microbial rain populations are provided in the graph. Red line is smoothing curve (LOESS). Grey line indicates relative abundance above 1 (log −4.6). OTU numbers are given in parenthesis and results from this analysis is further detailed in Table S2.

### Evaluation of metapopulation models in the Baltic Sea Proper

The relationship between local relative abundance and regional occupancy may be derived from and be directly linked to OTU abundance patterns. Analysis of distribution patterns among OTUs showed a recurring occupancy-frequency distribution that was significantly bimodal in nine out of eleven sampled months (Mitchell-Olds & Shaw’s test, p<0.01, n=6-12; Fig. 2). In these months, the bimodal pattern had one mode with most species found at a single site. This mode was followed by an initial monotonic decrease in the number of populations detected with increasing number of sites surveyed. We then noted a second mode with an increase in the number of populations that occurred at all sites. Thus, at a majority of sampling occasions, bimodal patterns were observed matching predictions of bimodality in the CSH model proposed by Hanski (16) rather than predictions of unimodality in Levin’s model (27). To our knowledge, our observed bimodal pattern represents the first finding of bimodality for prokaryotic communities as well as for organisms in marine environments in general.

**Figure 2.**
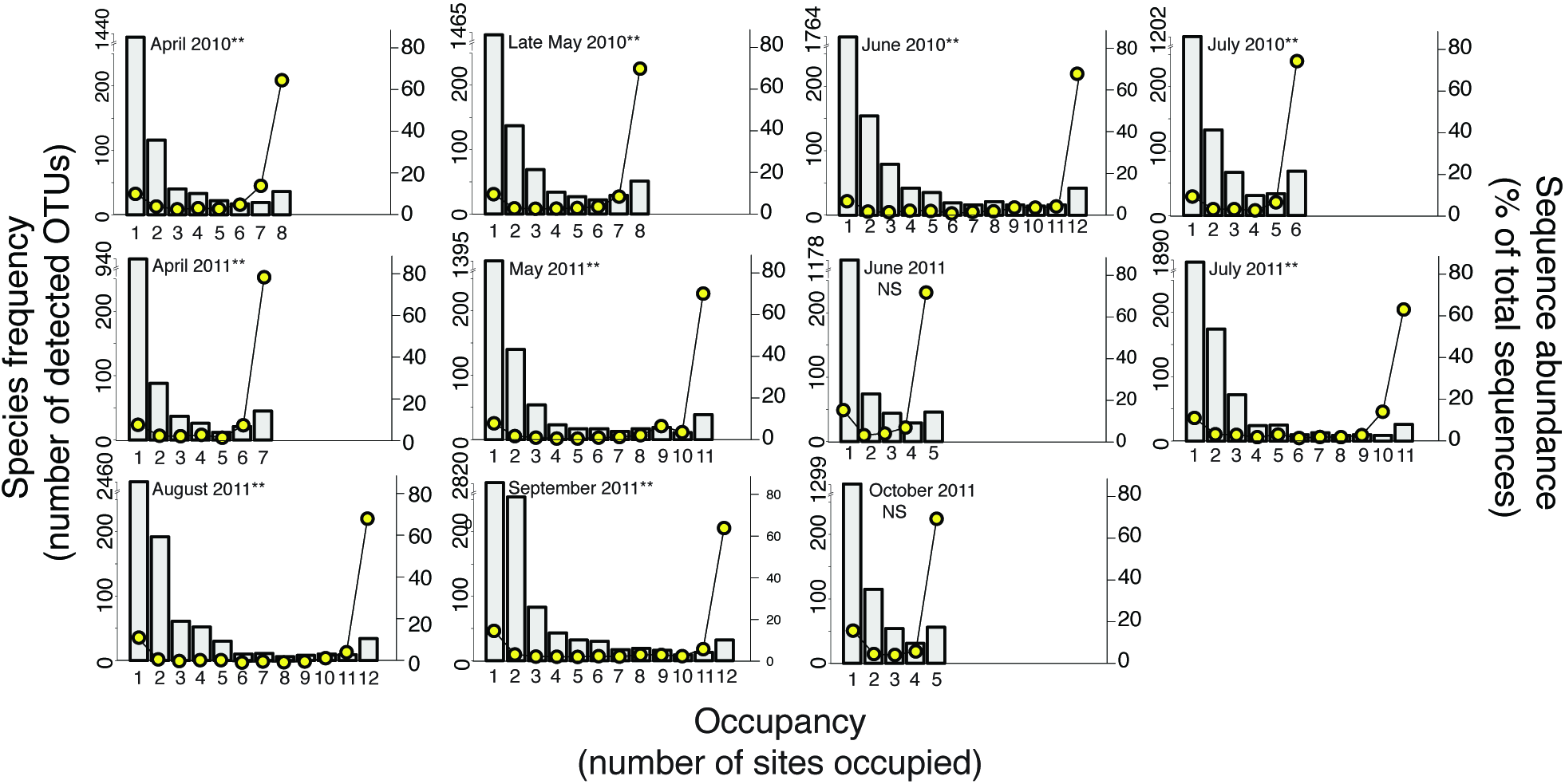
Occupancy-frequency distribution of OTUs in our spatiotemporal study of the Baltic Sea Proper (Western Gotland basin and Kalmar sound). Filled yellow circles indicate relative sequence abundance (percentage of total sequences in each sample). Asterisks denote significance levels (*P<0.05, **P<0.01, ***P<0.001) for Mitchell-Olds & Shaw’s test. NS denote non-significant bimodal patterns. The maximum number of sites sampled each month and at each station (i.e. maximum occupancy) is given by the x-axes.

Metapopulation models that include positive feedback mechanisms provide testable predictions of colonization and extinction rates (28). To elucidate how our observed data fit with current models, we calculated rates of colonization and extinction by comparing occupancies of OTUs between succeeding months, i.e. how many new sites an OTU had colonized compared to from how many previously occupied sites it had disappeared (Fig. 3; Table S1; Table S2). As in previous work (37), we considered the terms colonization and extinction in a broad sense as analogous to gain and loss of OTUs, i.e. local extinction is going from presence to absence. Accordingly, absence is indicated by local relative abundance <0.006%, given the detection limit of our sequencing methodology. In an analogous manner, colonization represents the shift from absence (i.e. below detection limit) to presence, irrespectively of whether this presence resulted from on-site growth of microbes from initially very low abundance or from immigration of populations from neighboring waters. Observed patterns in regional colonization rates followed a quadratic curve, where colonization (*C*) at first increased with fraction of sites occupied *(P)* until about 50-65% occupancy, followed by a decrease in *C* until maximum *P* (Fig. 3). Extinction rates followed a similar quadratic pattern, where extinction *(E)* first increased with *P*, but until a higher occupancy compared to *C* (between 65-75%), followed by a decrease of *E* until maximum *P* (Fig. 3). We examined how metapopulation models could explain variations in colonization and extinction rates by using non-linear least squares analysis. Predictions based on the CSH model proposed by Hanski (16) agreed well with the field results for extinction rates (See blue dashed lines in Fig. 3), with on average low residual standard error, whereas colonization rates had higher residual standard error (Table S1). The CSH model significantly explained both colonization and extinction rates. Predictions based on Levin’s model (27) showed overall higher residual standard error for extinction rates compared to the CSH model (Table S1; S2). Notably, both models significantly explained observed extinction rates, although extinction rates predicted by Levin’s model had lower parameter estimates compared to the CSH predictions (Table S1 and S2). Thus, both models produced significant fits for quadratic and linear colonization and extinction rates. However, since the CSH prediction of bimodal occupancy-frequency patterns matched our data on OTU distributions, whereas Levin’s predictions of unimodality did not, the CSH yielded an overall better agreement with the observed data. This agreement of theoretical predictions and measured bacterioplankton population dynamics favored the CSH model for interpreting regional effects of OTU distributions in the Baltic Sea Proper.

**Figure 3.**
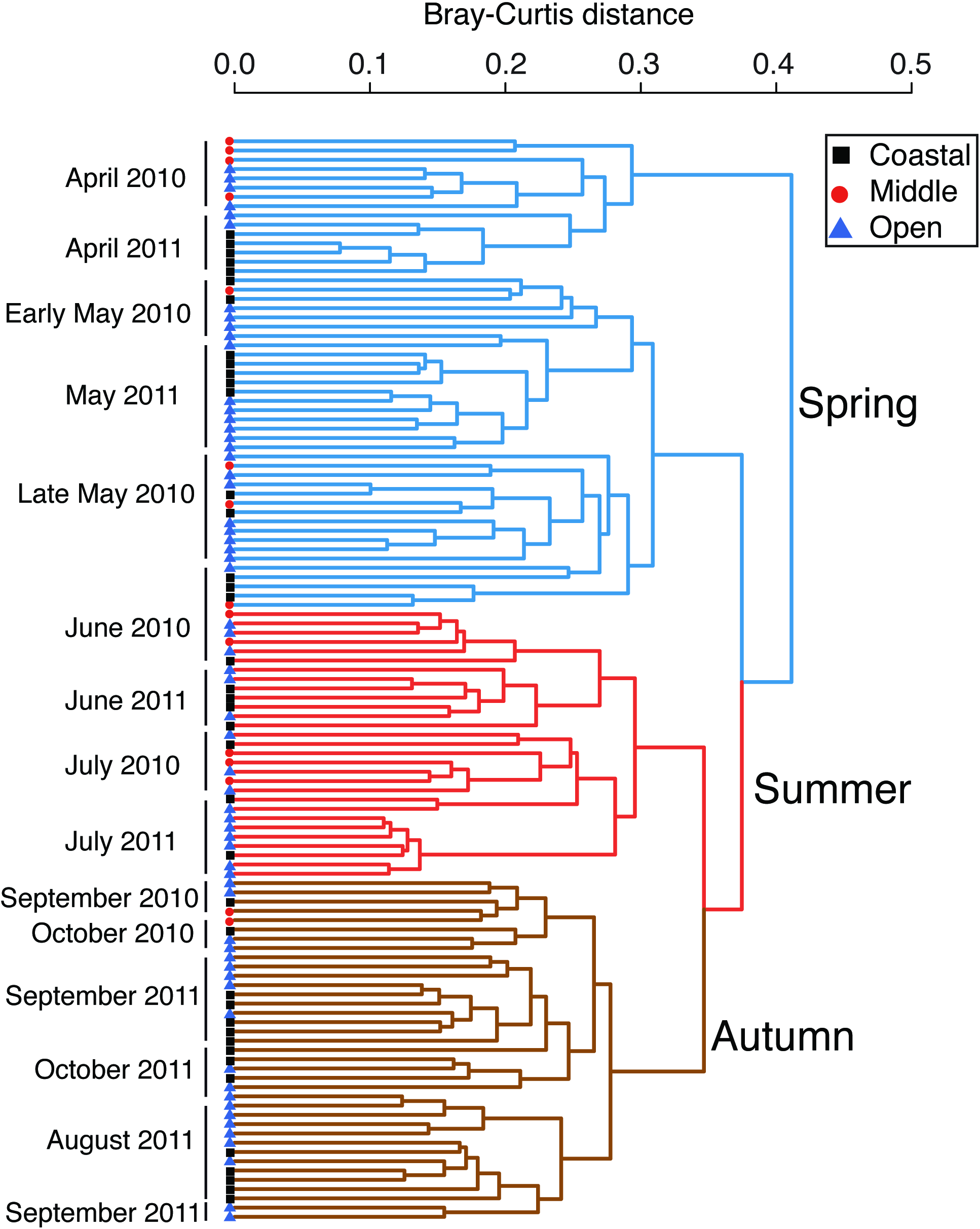
Colonization (*C*) and extinction *(E)* rates against occupancy *(P).* Colonization rates (A), and extinction rates (B). Each point represents the change in fraction of sites occupied from one month to the next for individual OTUs, in total 4437 OTUs for all months. The red line indicates mean and the red shaded area around the mean is standard error. Blue dashed and brown lines indicate the curve of observed data fitted by non-linear least squares to the metapopulation model by Hanski (1982) and Levin (1974), respectively, (Table S1 and Supporting information). Examples are from April, June and September from both years. OTU positions are jittered to reduce overplotting and colored in grey scale according to OTU density.

### Linking metapopulation dynamics and specific populations in the Baltic Sea Proper

Beyond the recognition of metapopulation models for estimating colonization and extinction rates of populations at a regional scale, the models function to identify ecological processes that are likely to be involved in determining the distribution of individual populations. In the Baltic Sea Proper dataset, the number of endemic satellite bacterioplankton populations (i.e. only found in one site each month) varied between 940 and 2820 OTUs. In contrast, the much fewer cosmopolitan core populations (i.e. found in all sites each month) ranged between 26 and 69 OTUs. A striking consequence of the steep rank abundance distribution patterns was that the OTUs found at all sites during one sampling month contributed to between 62 and 78% of the sequence abundance that month (see yellow filled circles in Fig. 4). Thus, although a grand majority of OTUs occupied a single sampling site each, these OTUs contributed only a very limited proportion of the total sequence abundance compared to the fairly few OTUs occupying all sites. It is therefore reasonable to assume that a community consists of a set of cosmopolitan core populations with high local and regional abundance compared to endemic satellite populations with low local and regional abundance. These findings reveal the importance of assessing core and satellite metapopulation dynamics to understand the genomic potential among bacterioplankton in biogeochemical processes such as carbon cycling. For example, in environmental metagenomic, transcriptomic or single-cell genomic approaches it is not possible to determine satellite or core characteristics per se within the data if samples were collected at a single local site. We infer that interpretations of such-omics approaches could be much aided by complementary analyses of 16S rRNA gene amplicon data at relevant spatio-temporal scales to couple regional dynamics of OTUs for assessment of satellite and core properties. Alternatively, conducting-omics approaches on many samples distributed over relevant spatiotemporal scales may be necessary to distinguish regional dynamics in microbial communities and interpretations thereof.

**Figure 4.**
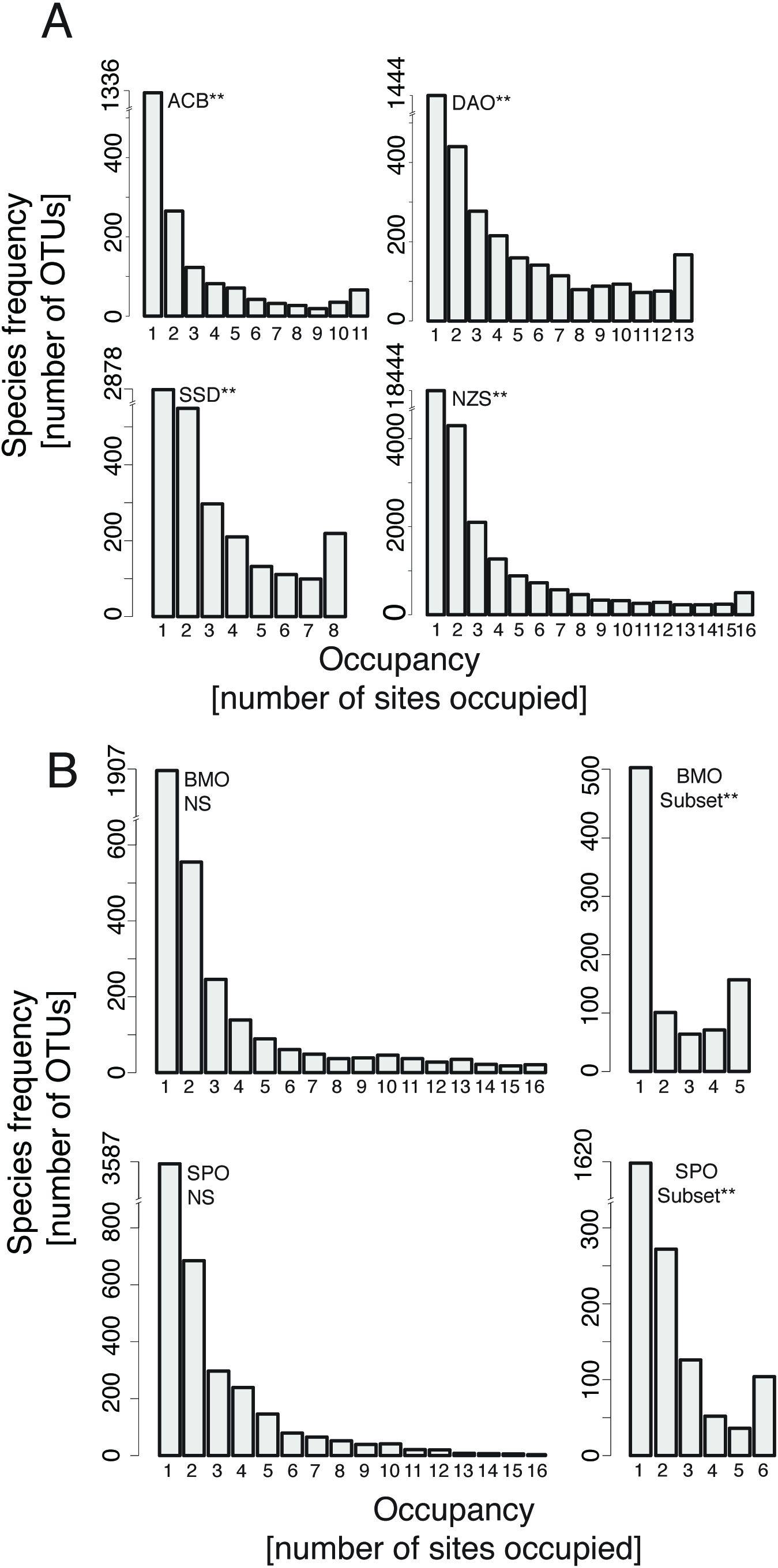
Examples of occupancy-frequency distribution of OTUs from global ICoMM datasets displaying significant bimodal patterns (A), examples not showing significant bimodality unless applying the analysis on a subset of samples within datasets (B). Oceanic regions are abbreviated; ACB - Arctic Chukchi Beaufort, DAO - Deep Arctic Ocean and NZS - New Zeeland Sediment, SSD - Spatial Scaling Diversity (Cape Cod Atlantic Ocean), BMO - Blanes Bay Microbial Observatory (North West Mediterranean Sea), SPO - Sponge Bacteria (Great Barrier Reef). Please note break in axes for the leftmost bar in each graph. Asterisks denote significance levels (*P<0.05, **P<0.01, ***P<0.001) for Mitchell-Olds & Shaw’s test. NS denote non-significant bimodal patterns.

In the present study we aimed to couple the identity of particular OTUs found in the analysis of local abundance and regional occupancy with observed metapopulation dynamics. Thus, populations were defined according to their local relative abundance (i.e. populations/OTUs >1% as abundant, <0.1% as rare (33), and 0.1-1% as common (38-40)). First, abundant populations with little variance in occupancy and relative abundance (SE <0.025 and <0.2%, respectively) were classified as “core”; corresponding to Hanski’s core populations, (n=6) (Fig. 1; Table S3). “Core” populations such as those belonging to the *Synechococcus* (PF_000001), CL500-29 (PF_000000), SAR11 (PF_000002), SAR86 (PF_000007), hgcI (PF_000008) and NS3a clades (PF_000009) were almost never found in the rare fraction in our study, and were at nearly all times part of the rightmost column in each of the panels in Figure 2. The presence of such cosmopolitan core populations is in agreement with the dominance of abundant lineages such as SAR11 and SAR86 clades typically found in marine environments (40-43). We have no immediate knowledge as to the factors that make such particular “core” populations successful. However, it is possible that “core” OTUs have advantages in utilization of widespread resources compared to other OTUs (17, 30). Hence, “core” OTUs may be linked to the concept of generalist dynamics (4, 7). Interestingly, for soil bacteria there is a significant coupling between habitat breadth and larger genome size, metabolic adaptability and occupancy, indicating advantages for versatile generalist populations (44). In the marine environment bacterial populations can also be widespread despite significant genome streamlining, e.g. the SAR11 and SAR86 lineages, suggesting that other mechanisms than genome size can shape microbial biogeography (45). In fact, many core populations found in the present study are predicted by phylogenetic affiliation to have streamlined genomes. We propose that positive feedback mechanisms between local abundance and regional occupancy can contribute to explain the widespread distribution of core OTUs, such as populations in the SAR11 and SAR86 clades and *Synechococcus*. This relative stability may be maintained despite potential local selection pressures by habitat filtering and predation/grazing. In contrast, stochastic variation in colonization and extinction rates has a large effect on populations with intermediate occupancy. Taken together, our data suggest that core OTUs depend largely on maintaining high levels of occupancy, implying an increased probability of colonizing new sites and sustaining large population sizes through positive feedback mechanisms.

Second, populations fluctuating between being abundant, rare and common with large variation in occupancy (SE >0.100) were classified as “transient” (n=38) (Fig. 1; Table S3). Interestingly, members of the *Cyclobacteriaceae* (PF_000047) and *Sporichthyaceae* (PF_000021) and the NS11-12 clade (PF_000030) mostly entered and exited between the abundant and common fractions of the bacterioplankton, and did not become rare. In contrast, the SAR92 (PF_000017) and *Spartobacteria* (PF_000011) OTUs fluctuated between the abundant and rare fractions.

Third, rare populations (<0.1% of bacteria) with little variance in relative abundance (SE < 0.2%), and high occupancy, were classified as “continuously rare” (n=9) (Fig. 1; Table S3). It is noteworthy that these continuously rare populations, e.g. CL500-29 (PF_000176), GOBB3-C201 (PF_000254), *Reichenbachiella* (PF_000068), *Thiobacillus* (PF_000187) and *Owenweeksia* (PF_000073), had a high occupancy. This potentially suggests that some populations had a permanent and widespread ecological strategy of being rare. In agreement, Galand and colleagues (46) observed how rare phylotypes remained rare when surface seawater samples were compared with deep waters from the Arctic, highlighting that most rare phylotypes were continuously rare within that ecosystem.

Fourth, rare populations with low occupancy were classified as “satellite”; equivalent to Hanski’s satellite populations, (n=4369) (Fig. 1; Table S3). Most “satellite” populations were only found in a single site, e.g. *Sulfurimonas, Thermoplasmatales* (PF_012272), SAR11 (PF_002803), SAR86 (PF_003406), *Brevundimonas* (PF_001083) and *Pedobacter* (PF_008706). The satellite characteristics of rare populations indicate a clear endemic biogeography of the rare biosphere, likely caused by the high extinction rates observed. In accordance, the rare biosphere has a distinct biogeography despite the potential for high dispersal and low loss rates (46). Thus, in contrast to core populations, rare and geographically more restricted satellite populations will be less likely to colonize new sites and more likely to become locally extinct.

We note that the observed extinction rates were always higher than colonization rates (>12.5 and <12.5, respectively) and that the parameter estimates of extinction and colonization rates predicated by the core-satellite model were significantly different (Fig. 3; Table S1; two-sample t-test, p=4.02x10^−11^, n=21). Higher rates of extinction compared to colonization imply that a few populations with higher colonization compared to extinction rates cause bimodal occupancy-frequency patterns. However, most populations are headed towards local extinction and regional rarity. If a few core populations did not exhibit higher colonization rates than extinction rates through positive feedback mechanism, most populations would eventually disappear. The quadratic relationships for colonization and extinction rates at different occupancies highlight the importance of regional dynamics for local populations with a decreasing probability for extinction with increasing occupancy. Overall, these results substantiate the hypothesis that the effect of stochastic variation is highest for populations with intermediate occupancy. A corollary of this scenario is that it is far more likely for common populations than for rare ones to become more common, and more likely for rare populations to become even more rare. Hence, we validated the core-satellite hypothesis, showing that positive feedback mechanisms regulate bacterioplankton population dynamics in the Baltic Sea Proper.

In our Baltic Sea Proper dataset we also observed special cases of population dynamics characterized by high colonization or extinction events. Firstly, several OTUs showed a pattern of being absent one month, and present at nearly all sites the following month, thus having a large colonization rate (Fig. 3). We denote this special case of colonization dynamics as “microbial rain” pattern, as a reformulation of the concept "propagule rain" coined by Gotelli et al. (29) for dispersal patterns of seed propagules from trees. The detected “microbial rain” populations, i.e. with very large variation in occupancy, included *Roseobacter* (PF_000153), *Fluviicola* (PF_000171) and *Brevundimonas* (PF_000126). A reciprocal pattern was observed for extinction rates, where some OTUs were present at all sites one month, next to be absent from all sites the following month - we term this “microbial evanescence”, indicating large extinction rates. “Microbial evanescence” populations were exemplified by *Owenweeksia* (PF_000313) and two *Synechococcus* OTUs (PF_000143, PF_000060). We suggest that “microbial rain” and “microbial evanescence” types of OTU distributions could result from linkages between strong seasonal shifts in environmental conditions and niche differentiation (7). Alternatively, wind-driven upwellings or other hydrological events may also cause rapid shifts in bacterial community composition (47, 48). Still, such hydrological events could not have been common in our study system, considering the relatively stable seasonal succession and smaller impact of spatial differences over the two studied years (Fig. S2). Similarly, analysis bacterioplankton population dynamics in an adjacent Baltic Sea Proper sampling site showed distinct seasonal succession related to environmental conditions such as temperature, but only occasionally influenced directly by hydrological disturbances (38).

Altogether, we show that the core-satellite hypothesis can be applied to prokaryotic assemblages and be used to characterize biogeographical patterns providing insight into the importance of regional dynamics. It is noteworthy that the core-satellite hypothesis also provides a mechanistic understanding of the division between rare and abundant bacterioplankton populations in the local environment. Thus, the core-satellite hypothesis can be used to interpret the common observation of log-normal OTU distribution patterns potentially caused by stochastic variation in colonization and extinction rates.

#### OTU abundance distribution in a Baltic Sea - Kattegatt transect

To extend the exploration of the applicability of the CSH and its consequences for interpreting bacterioplankton distribution patterns, we investigated complementary datasets that covered a range of marine environments and geographic distances. By applying our analysis to a separate dataset of 16S rRNA genes from the Baltic Sea we could assess bacterioplankton distribution patterns over the entire 2000 km salinity gradient of this semi-enclosed brackish water system (Fig. S4A; 19 sites). We did not find a significant bimodal occupancy-frequency pattern among surface samples when including marine sites from the Kattegat Sea (Fig. S4B; Mitchell-Olds & Shaw’s test, *p*>0.01, n=19). However, when excluding the marine sites from the brackish water sites we found a significant bimodal pattern (Fig. S4C; Mitchell-Olds & Shaw’s test, *p*<0.01, n=17). The differentiation between marine and brackish water sites indicated that changes in environmental conditions, in this case likely a salinity shift (from 5.62 ± 2.06 to 22.00 ± 3.10, Baltic Sea and Kattegat Sea respectively), potentially influenced the metapopulation dynamics and separated bacterioplankton assemblages into distinct biomes in the Baltic Sea compared to the Kattegat Sea. In fact, salinity regulates the distribution of bacterial populations and their functional potential in the Baltic Sea (49, 50), and was recently suggested to constitute part of a global brackish microbiome (10). We also note that strong shifts in other marine biota occur at the border between brackish-marine conditions, also known as the Darss Sill, as shown in e.g. palaeoecological analyses of diatoms (51). We conclude that Baltic Sea bacterioplankton communities exhibit core and satellite population dynamics as observed in the Baltic Sea Proper data. Thus, species-abundance patterns among Baltic Sea bacterioplankton may provide clues to how bimodality is linked with specific biomes.

#### OTU abundance distribution in global ICoMM datasets

To further extend the analyses of bimodal occupancy-frequency distributions, we carried out analyses of distribution patterns on broad global biogeographic datasets provided in the frame of ICoMM (52). This showed that bacterial communities in 17 of 31 oceanic regions displayed a significant bimodal pattern (Table 1). In Fig. 4A we exemplify some of the statistically significant bimodal patterns by the samples from the Arctic Chukchi and Beaufort Seas (ACB), Deep Arctic Ocean (DAO), New Zealand sediments (NZS) and open ocean off Cape Cod (SSD) (Mitchell-Olds & Shaw’s test, *p*<0.01, n=8-16; Fig. 4A; Table 1). It is noteworthy that bacterial community composition in several vast oceanic regions (e.g. ACB and NZS) displayed bimodality. Still, in datasets from several regions, OTU occupancy-frequency distribution patterns were not significantly bimodal (Table 1), which can be exemplified by the data sets from the northwest Mediterranean (BMO) and Sponge communities from the Great Barrier Reef (SPO) (Mitchell-Olds & Shaw’s test, *p*>0.01, n=11-16; Fig. 4B; Table 1). However, some of these datasets stand out by the heterogeneity of the samples they encompass. For example, it is well established that bacterioplankton community composition differs significantly between depths in a range of ecosystems, including the northwest Mediterranean Sea (see e.g. (53). Still, the BMO transect comprised samples from different depths. We therefore separated samples according to depth and analyzed surface waters separately; we thus found a significant bimodal occupancy-frequency pattern for these surface samples (too few samples were available to test bimodality at other depths). In a similar manner, it is established that bacterial community composition differs considerably between different sponge species (54). Thus, when accounting for sponge species, we found a significant bimodal occupancy-frequency pattern for OTUs associated with the sponge *Rhopaloeides odorabile* in the SPO dataset. Taken together, analyses of datasets from several oceanic regions provided support for the widespread prevalence of bimodality among marine prokaryotic assemblages.

**Table 1.**
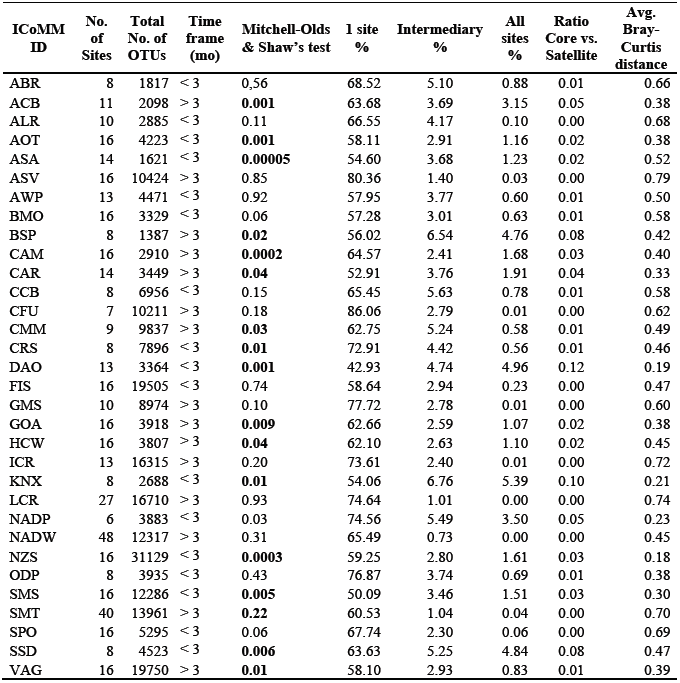
Bimodality in global ICoMM data of marine microbes. Regions with >5 spatially different sites (horizontal or vertical) were selected. For each ICoMM dataset, we have included the number of sites, total number of OTUs and the time frame within which the samples were collected, with p-values for Mitchell-Olds & Shaw’s test of quadratic extremes. We further provide percentages of OTUs occupying 1 site, intermediary sites (i.e. average percentage of OTUs occupying 2 or more sites but not all) and all sites. The ratio between core (i.e. OTUs occupying all sites) and satellite (i.e. OTUs occupying 1 site) populations is also displayed. For comparison, average Bray-Curtis distances for each dataset is also provided.

From the above we deduce that colonization of bacterioplankton populations could be regulated by physicochemical factors, thus structuring community composition into specific biomes. In such a scenario, large differences in environmental conditions resulting from e.g. differences in depth, salinity or nutrient concentrations and/or physical oceanography within or between oceanic regions should lead to distinct patterns of bacterioplankton community structure, i.e. no bimodality. In contrast, regions within which dispersal is high compared to local environmental conditions bimodality should be detected. We therefore evaluated the relationship between bimodal occupancy-frequency patterns and beta-diversity and environmental conditions. To analyze such potential dependencies, we calculated average Bray-Curtis distances for all ICoMM datasets and separated these distances into two groups for datasets with significant or non-significant bimodal patterns, respectively. This revealed a significant difference between groups with and without significant bimodal patterns (two-sample t-test, p=8.09x10^−7^, n=31); datasets with significant bimodal patterns on average had lower Bray-Curtis dissimilarity (Fig. 5A). In contrast, datasets with higher average Bray-Curtis dissimilarity were linked to non-significant bimodal patterns. Moreover, when comparing the ratio between core and satellite populations (percentage of OTUs occupying all sites compared to a single site) and Bray-Curtis dissimilarities, the ratio was significantly negatively correlated with average Bray-Curtis distance (linear regression, p=6.42x10^−4^, R^2^=0.42, n=31; Fig. 5B). A second-degree polynomial curve fit indicated a saturation where the ratio leveled off at around 0.04 (p=1.4x10^−5^, n=31; Fig. 5B). Similarly, the ratio between core and satellite populations was on average 0.05±0.03 in the Baltic Sea Proper dataset and monthly data coupled with average Bray-Curtis distances followed the negative slope (see open circles Fig. 5B). In terms of absolute changes in environmental conditions - i.e. analyses made on average Euclidean distances - temperature, salinity and depth had an overall tendency toward lower absolute differences for datasets with significant bimodal patterns (Fig. 5C-E). However, only depth (two-sample t-test, p=0.045, n=24) and temperature (two-sample t-test, p=0.036, n=20) were significantly different between datasets with and without significant bimodal patterns. Notably, absolute shifts in geographic distance were not significantly different between datasets with or without significantly bimodal patterns (Fig. 5F; two-sample t-test, p=0.292, n=26). This suggests that shifts in environmental conditions, and not spatial distance *per se*, are critical for regulating positive feedback mechanisms between local abundance and regional occupancy, thus affecting the occupancy-frequency pattern. Collectively, these data and changes in occupancy-frequency patterns may be highly valuable for distinguishing where shifts in biomes occur across environmental gradients.

**Figure 5.**
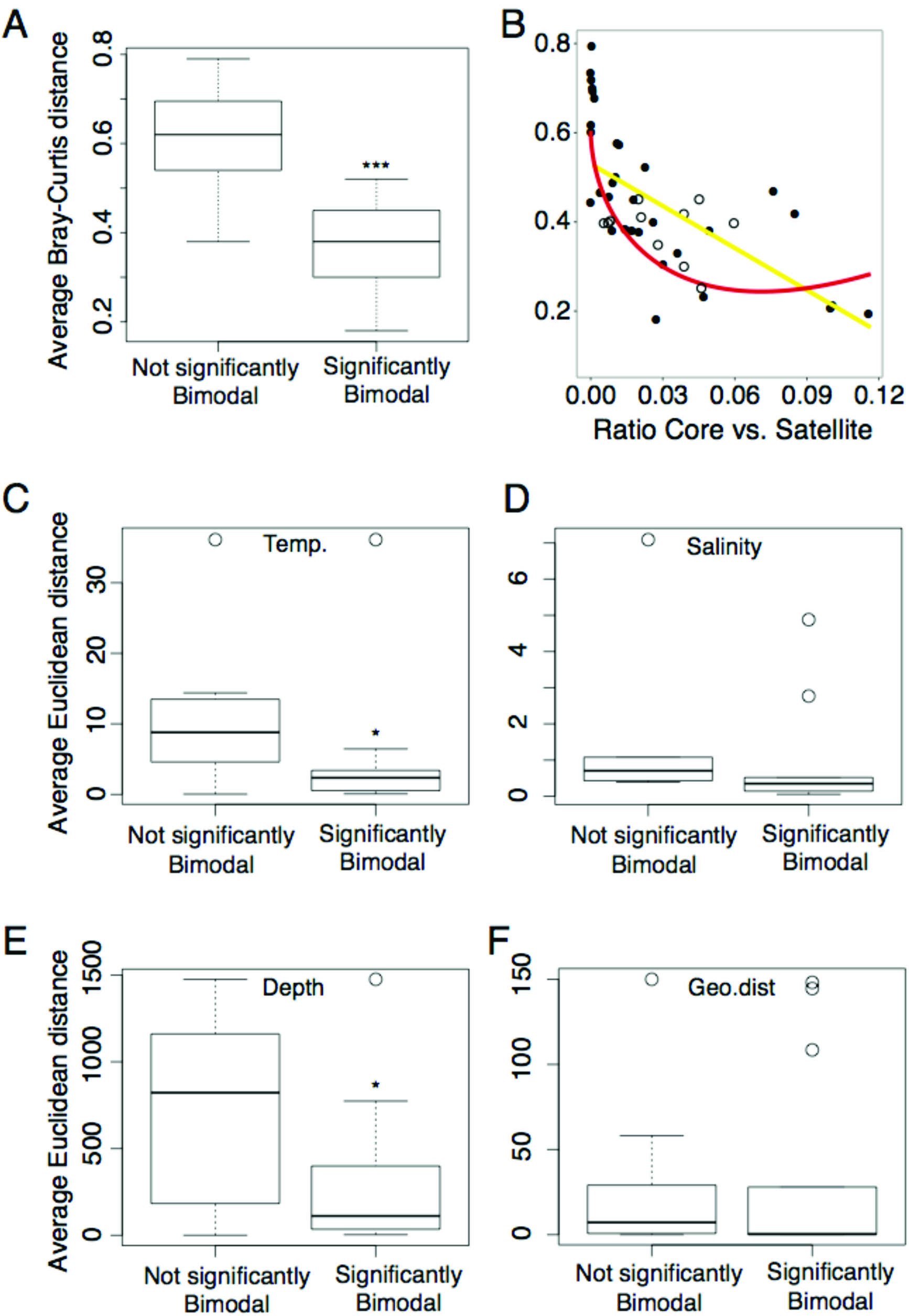
Boxplot of average Bray-Curtis distances from each ICoMM dataset and nonsignificant or significant bimodality (A), the relationship between average Bray-Curtis distance and the ratio between core and satellite populations (percentage of OTUs occupying all compared to one site) (B), and the relationship between average Euclidean distances of environmental variables, geographic distance and non-significant or significant bimodality (C-F). Open circles in (B) denote monthly Baltic Sea Proper samples collected in this study that were added independently of the regression and polynomial analyses. Asterisks denote significance levels (*P<0.05, **P<0.01, ***P<0.001) for two sample t-tests.

We are aware of only one study that has found bimodality in aquatic systems, and that study concerned fish in Amazonian lakes (55). The CSH has also been tested in a set of boreal stream systems, where it was found that diatoms did not exhibit bimodality (32). The discovery in the present study of bimodal distribution patterns, linking local abundance and regional occupancy of marine planktonic bacteria, may seem surprising, especially given the common perception that pelagic marine habitats are relatively homogeneous and allow efficient dispersal of prokaryotic assemblages. Nevertheless, differential distribution of marine bacterioplankton populations is frequently found to be linked with shifts in environmental conditions, such as salinity, temperature, phytoplankton biomass and DOC (25, 49, 56, 57). Interestingly, it has been suggested that due to the nature of free-living and easily dispersible prokaryotes, bimodal occupancy-frequency patterns should be relatively common (31). It is also noteworthy that explorations of occupancy-frequency patterns in highly diverse communities is lacking; typically <100 species are considered, whereas only a few have investigated communities with >400 species (28, 32). Thus, in the present paper we show for the first time linkages between highly diverse prokaryotic communities in marine environments and metapopulation dynamics.

Although the bimodality patterns observed in the current study conforms with Hanski’s metapopulation model, alternative hypotheses have been suggested to explain the shape of different occupancy-frequency distributions, including for example sampling effects (28, 29, 58). Still, in the datasets studied here, bimodality was found over a variety of temporal scales and over a diversity of geographical distances, indicating a pronounced robustness in this occupancy-frequency pattern. Alternatively, neutral mechanisms in combination with dispersal could shape a distribution pattern with locally abundant taxa also being frequently occurring (35, 36, 59). However, colonization and extinction rates predicted by the CSH agreed well with our field data and point toward that stochastic variation in rates of local extinction and/or colonization can account for observed bimodality. Taken together, our findings on the biogeography of microbial populations at distances ranging from <10 km to >10,000 km indicate that bimodal OTU distributions are an important recurring pattern in several marine ecosystems across the world’s oceans, stressing the importance of regional dynamics.

### Conclusions

It is becoming increasingly clear that community-structuring processes acting on macroorganisms also determine distribution patterns of marine bacterioplankton populations (60-62). Our extensive analyses suggest that, as generally recognized in terrestrial ecology, positive feedback mechanisms between local abundance and regional occupancy are important drivers of bacterioplankton community composition. As such, our findings demonstrate that metapopulation theory, developed for terrestrial ecology, also provides a tool for interpreting patterns of microbial biogeography. The application of the core-satellite hypothesis on prokaryotic assemblages in the sea forms a framework for interpreting biogeographical patterns and defining the division between rare and abundant bacterioplankton populations in the local environment. Characterization of metapopulation dynamics in marine bacterioplankton communities would not only allow for comprehensive biogeographical analyses of marine microbes, but also contribute to improve the definition of cosmopolitan and endemic populations and the mechanisms by which they are influenced. Our findings suggest that shifts in microbial biogeography, in particular microbial biomes as defined by bimodal occupancy-frequency patterns compared to unimodal patterns, may be used to describe and pinpoint changes in the environment likely caused by dispersal limitation for use in molecular biomonitoring.

## Material and Methods

### Field sampling

In 2010-2011 (April-October), spatio-temporal dynamics of bacterioplankton communities were investigated at 16 stations off the east coast of Sweden in the Baltic Sea Proper. The distance between stations was typically less than 10 km (Fig. S1). The study area was chosen as a part of collaboration between the projects PLANFISH and EcoChange to study food-web dynamics in a coastal to open ocean gradient. For details on hydrological conditions in the study area please see (63, 64). Seawater from each station was collected in acid washed Milli-Q rinsed polycarbonate bottles, at discrete depths (2, 4, 6, 8 and 10 m) that were pooled and filtered shipboard. All 16 stations were represented in 2010 and classified according to their spatial distribution (coastal-offshore) as coastal, middle and open sea stations. During 2011, only coastal and open stations were represented. Not all stations could be included every month, and we used instead a subset of 5-12 stations (in total 117 stations).

### DNA extraction, 454 PCR, sequence processing and analysis

Collection of DNA from the 117 samples was carried out by filtering 1-2 liter of seawater onto 0.2 μm pore size, 47-mm diameter Supor filters (PALL Life Sciences). The filters were immediately frozen at −80 °C in 1.8ml TE buffer (10 mM Tris, 1 mM EDTA, pH 8.0) until further processing. DNA extraction was performed according to the phenol-chloroform protocol in Riemann et al. (65). Bacterial 16S rRNA gene fragments was amplified with bacterial primers 341F and 805R (V3-V4 hypervariable region) containing adapters and barcodes following the protocol of Herlemann et al. (49). The resulting purified barcoded amplicons were normalized in equimolar amounts and sequenced on a Roche GS-FLX 454 automated pyrosequencer (Roche Applied Science, Branford, CT, USA) at SciLifeLab, Stockholm, Sweden. Raw sequence data were processed following the bioinformatical pipeline in Lindh et al. (66). Briefly, sequences from all samples were clustered together into operational taxonomic units (OTU) at the 97% identity level (sequence similarity) using USEARCH (67) and taxonomically identified using the SINA/SILVA database. The 454 runs resulted in 300,000 reads (2010) and 396,000 reads (2011). After denoising and chimera removal, samples contained on average 4176 (±1716 SD) sequence reads (2010) and 4432 (±1445 SD) sequence reads (2011) for each sample. The final OTU table consisted of 4437 different OTUs (excluding singletons). For alpha-diversity measures we subsampled the sequences to 2500 reads per sample. Rarefaction curves are provided in Figure S5. DNA sequences have been deposited in the National Center for Biotechnology Information (NCBI) Sequence Read Archive under accession number SRP023607.

As for the Baltic Sea Proper dataset surface water during the full Baltic Sea transect was collected on 0.2 μm pore size, 47-mm diameter Supor filters (PALL Life Sciences) and 16S rRNA genes were amplified with 341F-805R. Sequencing of the Baltic Sea transect were performed on Illumina MiSeq. Sequences were clustered into OTUs at 99% identity level using USEARCH (67). For the Baltic Sea dataset we subsampled the sequences to 10,000 reads per sample.

OTU tables (97% 16S rRNA gene fragment sequence identity) of bacterial sequences (V6 hypervariable region) from ICoMM projects were downloaded from the Visualization and Analysis of Microbial Population Structures (VAMPS) database (http://vamps.mbl.edu/index.php). ICoMM datasets were subsampled at 10,000 reads per sample. A map of the ICoMM stations is available at http://vamps.mbl.edu/google_earth/ge_icomm.php showing the location and spatial scale of the oceanic regions included in the analyses performed in the present paper.

### Statistical analyses and graphical outputs

All statistical tests were performed in R 3.2.2 (68), using the package Vegan (69). An equivalent for Tokeshi’s test of bimodality was performed using Mitchell-Olds & Shaw’s test for the location of quadratic extremes. Nonlinear least squares analysis was performed using the equations of Levin’s model and the CSH hypothesis. Levin’s original model (27) is calculated as follows:

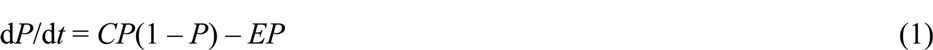

where *P* is the fraction of occupied sites, *C* is colonization rate and *E* is extinction rate. When *P* = 1, occupancy is 100% and all sites are occupied and when *P* = 0, occupancy is 0%, meaning regional extinction. Colonization rate is the number of colonized empty sites over time. If *C* is greater than *E* plus the variance in *E (C > E+ S^2^E)*, then Levin’s model predicts a unimodal distribution. Hanski’s model (16) is calculated as follows:

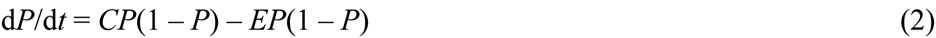

Thus, if the variance in *S (C - E)* is greater than *S* (i.e. σ^2^_s_ > *S*) the model predicts a bimodal species distribution.

Graphical outputs was made in R 3.1.2 using the package ggplot2 (70). Figures S1 and S4A were made using ODV (Version 4.5.0).

## ACKNOWLEDGMENTS

We acknowledge Sabina Arnautovic and Emmelie Nilsson for their skillful technical assistance in the processing of samples. Emil Fridolfsson at the Linnaeus University Centre for Ecology and Evolution in Microbial model Systems (EEMiS), Olof Lövgren at the Swedish University of Agricultural Sciences (SLU) and the crew of R/V Mimer for their outstanding effort in this sampling campaign. We also thank Bengt Karlson at the Swedish Meteorological and Hydrological Institute (SMHI), for the sampling of the trans-Baltic bacterioplankton communities on R/V Poseidon. This work was supported by the Swedish Research Council and the Swedish governmental strong research programme EcoChange and the Linnaeus University Centre for Ecology and Evolution in Microbial model Systems (EEMiS). Further support was provided by the PLAN FISH project, financed by the Swedish Agency for Marine and Water Management (former Swedish Board of Fisheries) and the Swedish Environmental Protection Agency.

## Supplementary Figures and Tables

**Figure S1.**
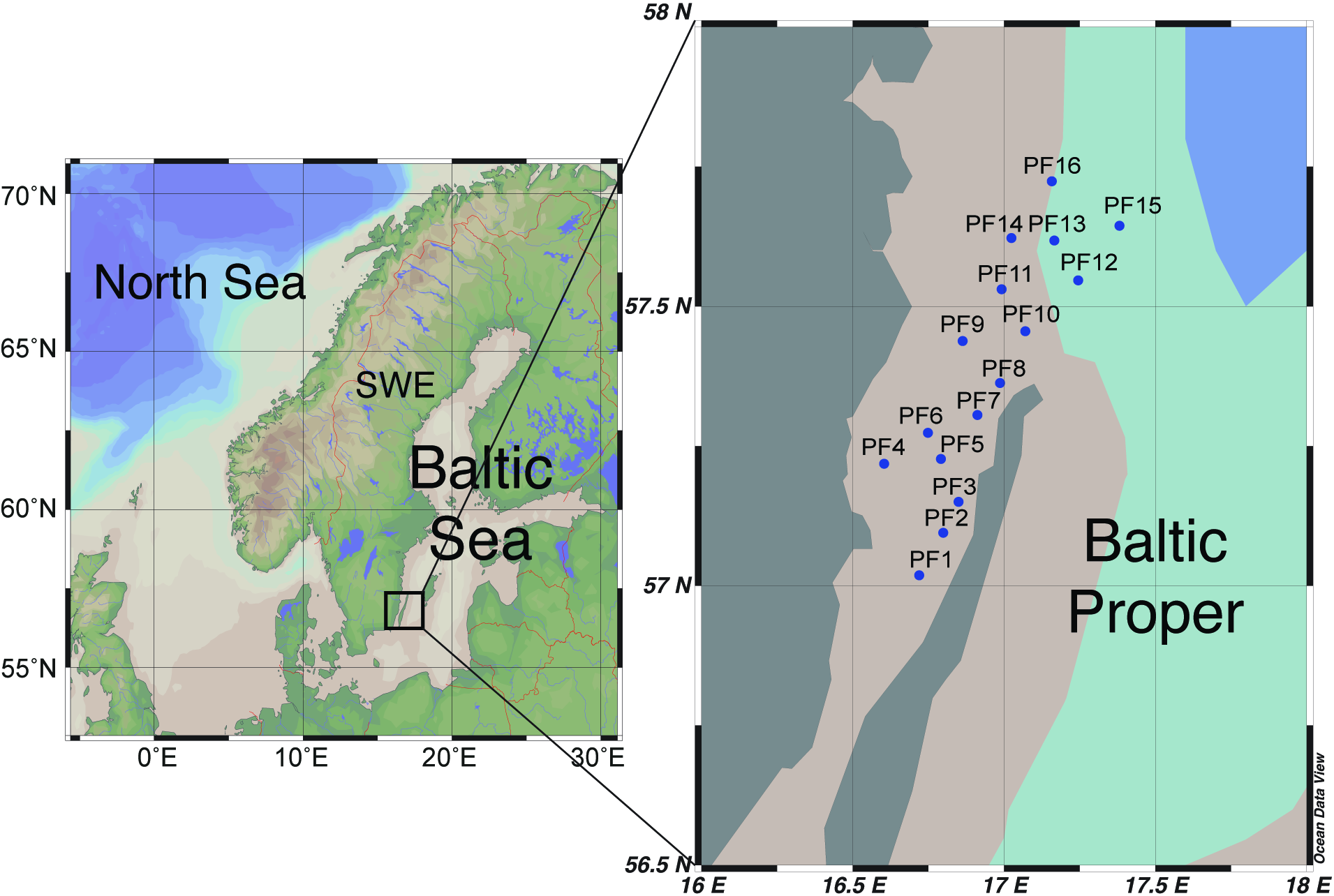
Map showing the 80-km transect in the Baltic Sea Proper, adapted from Legrand et al. (63, 64). This transect was performed in the PLANFISH and EcoChange frameworks, (see e.g. (63, 64)).

**Figure S2.**
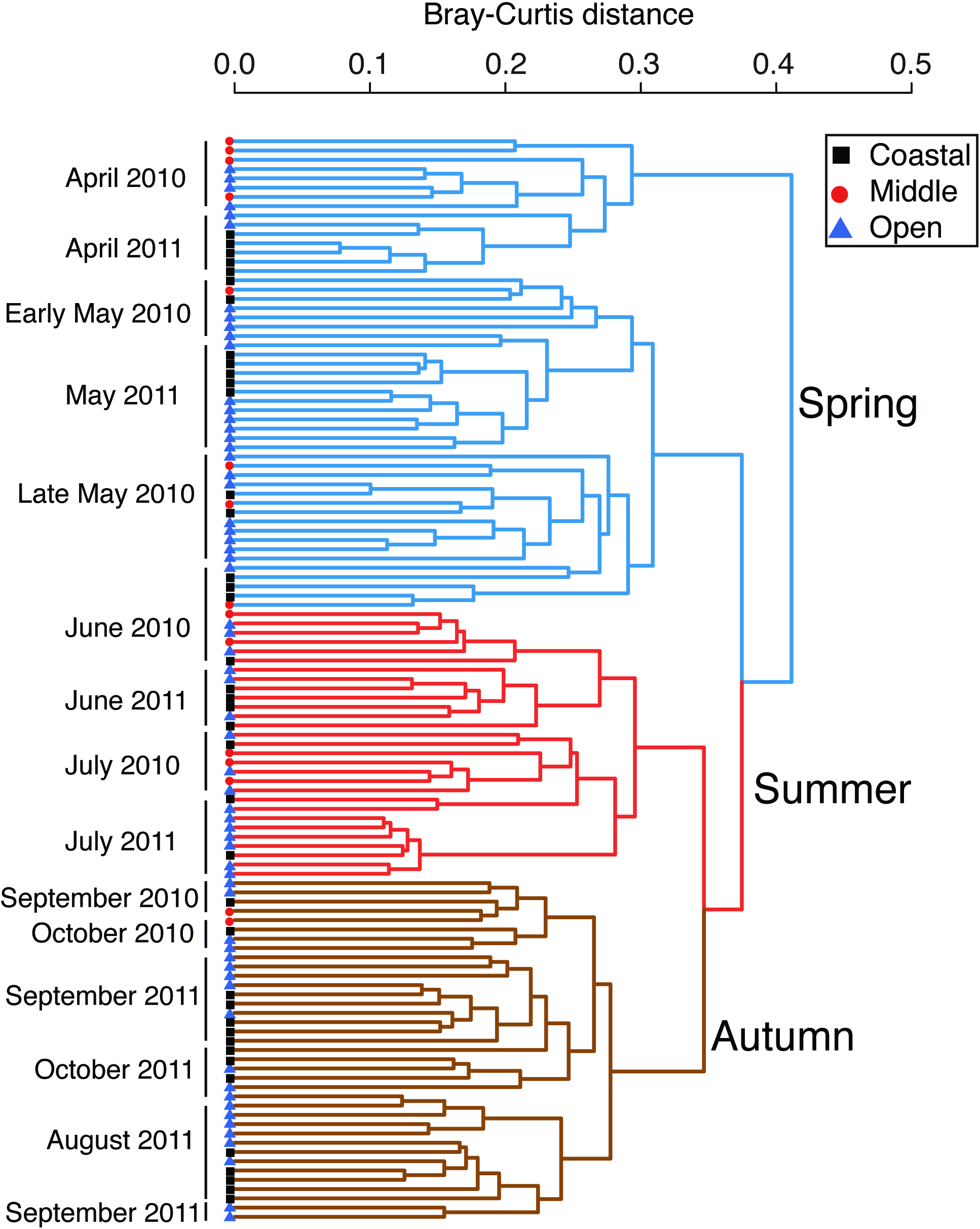
Cluster analysis for comparing beta diversity calculated from Bray-Curtis distance estimation based on 454 pyrosequencing data using 97% 16S rRNA sequence similarity. Color of clusters indicates seasonal communities. Symbols indicate coastal, middle or open ocean sites.

**Figure S3.**
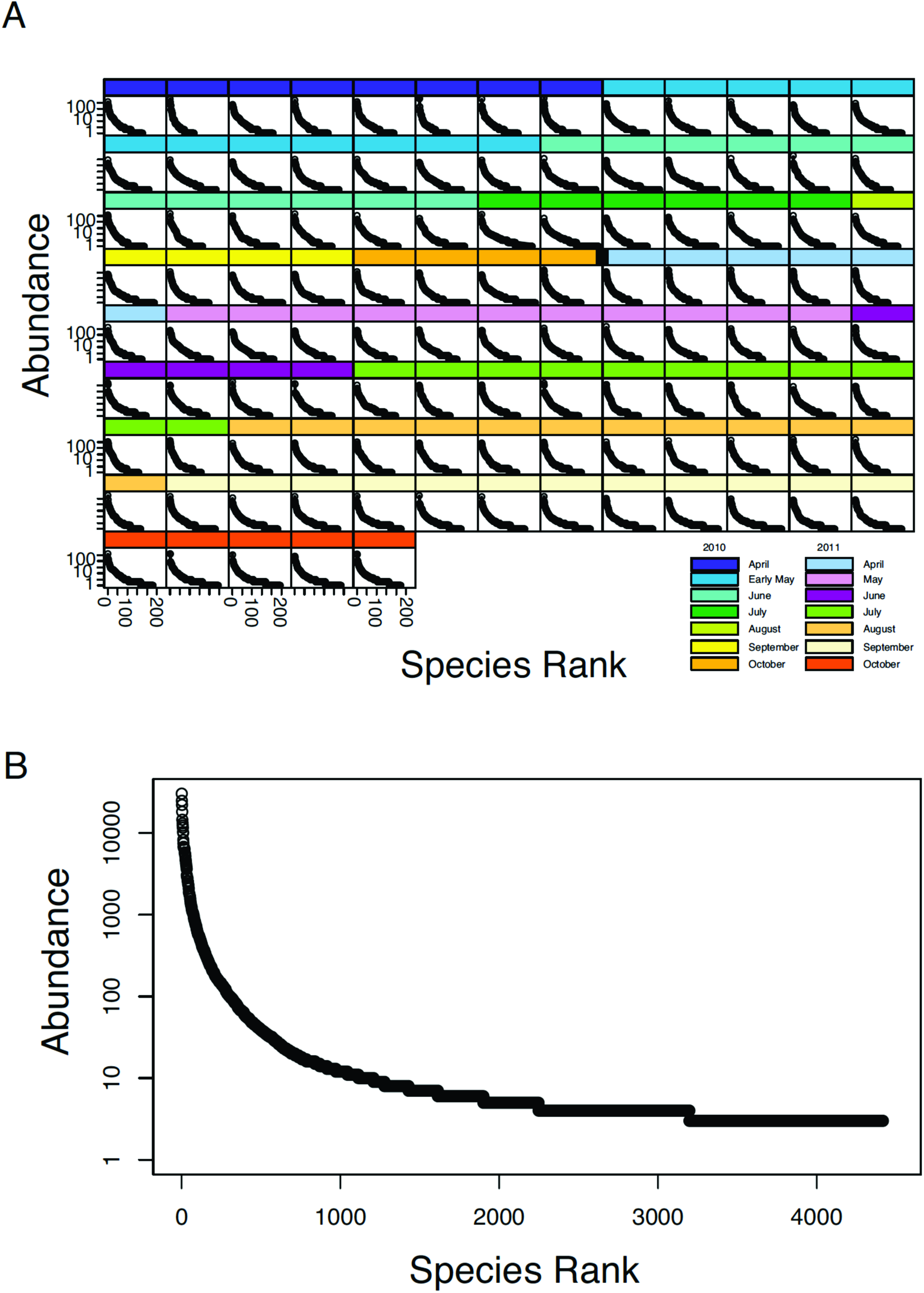
Baltic Sea Proper species rank-abundance curves for separate samples (A), and for all samples collectively (B). All were log-normal, with a few dominant OTUs and a tail of rare OTUs.

**Figure S4.**
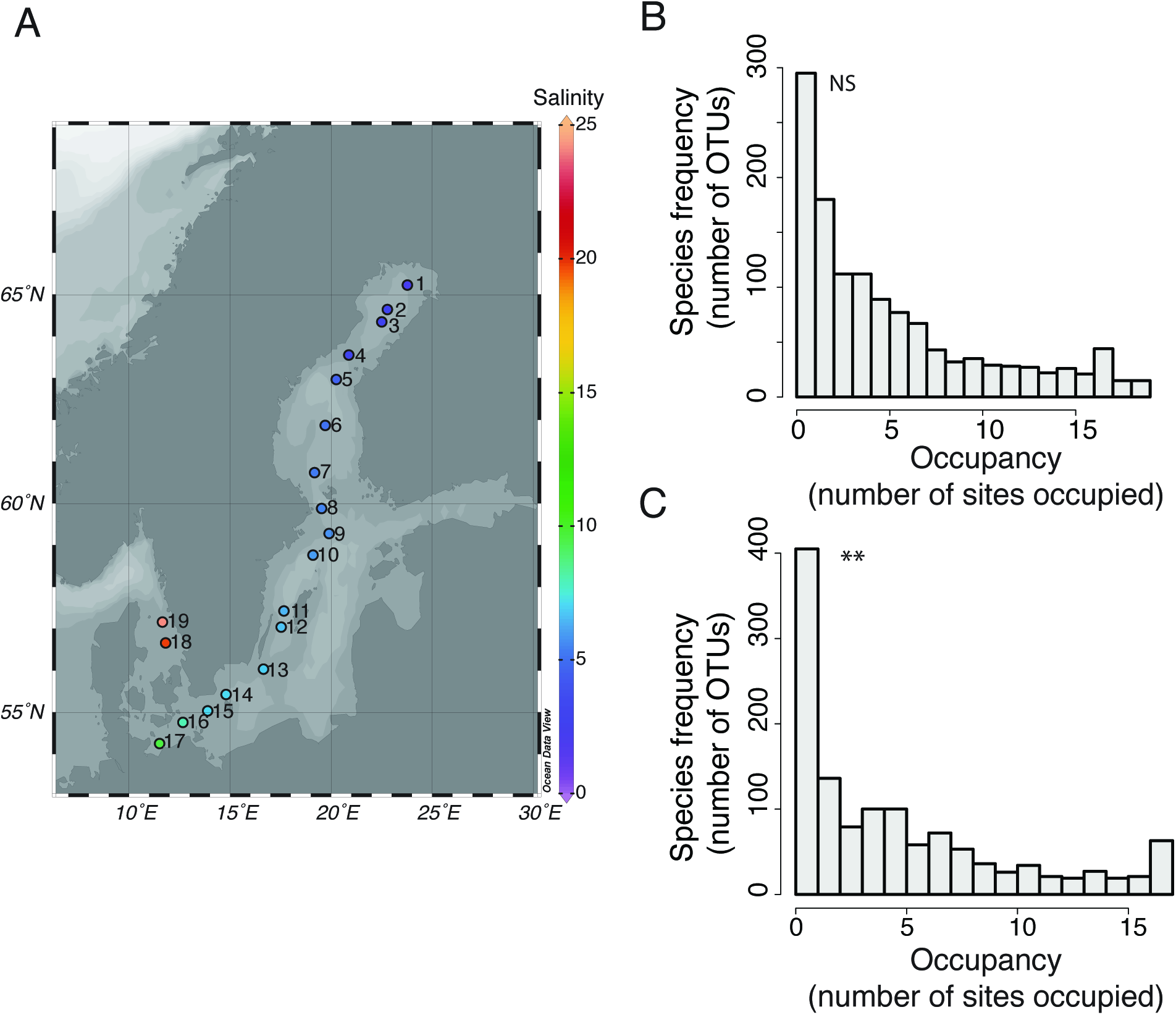
A Baltic Sea transect covering the entire 2000 km salinity gradient (A), with the occupancy-frequency distribution of OTUs for all sites of the transect (B), and the occupancy-frequency distribution of OTUs for brackish water sites only (i.e. salinity <15). Asterisks denote significance levels (*P<0.05, **P<0.01, ***P<0.001) for Mitchell-Olds & Shaw’s test. NS denote non-significant bimodal patterns.

**Figure S5.**
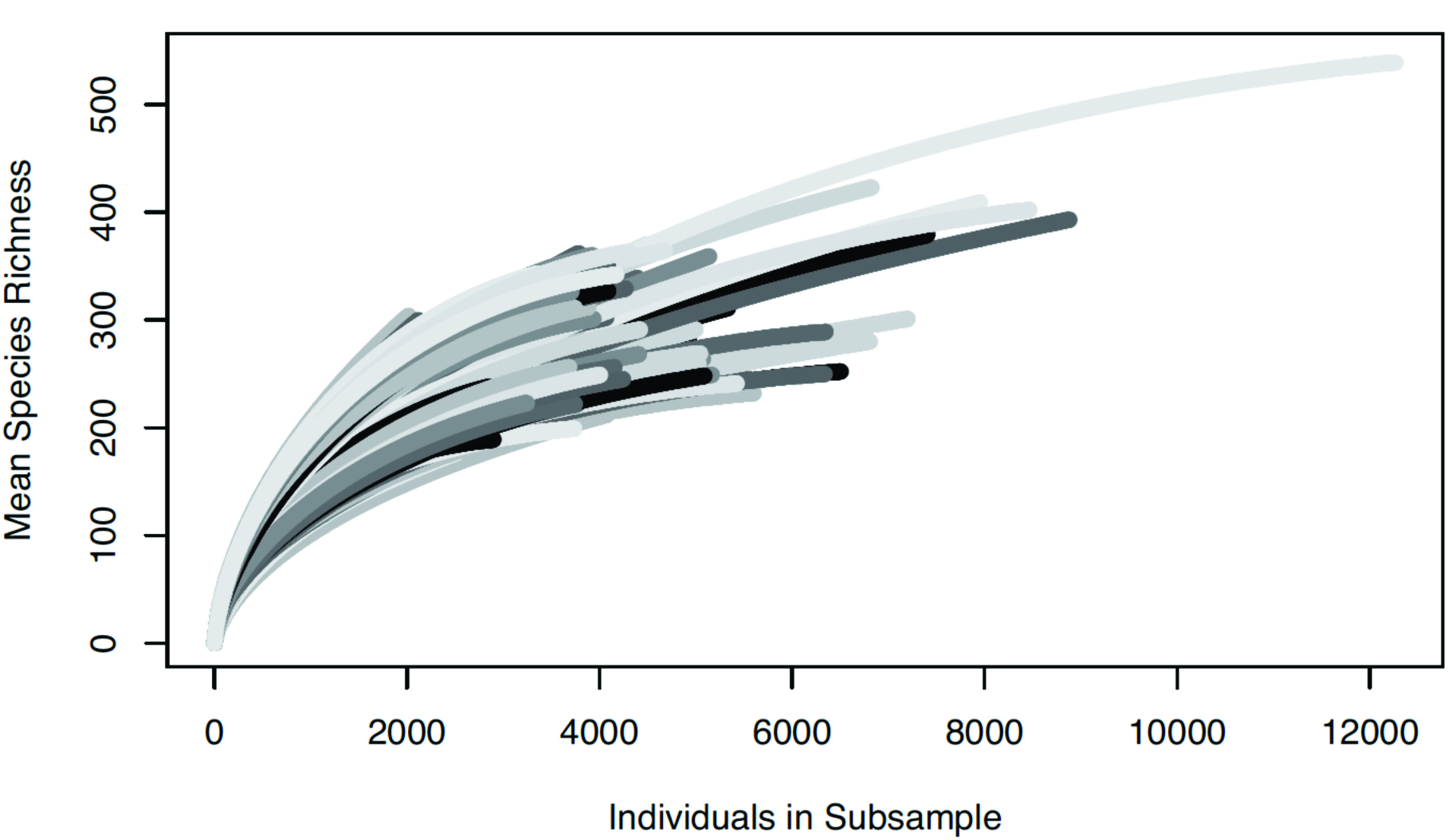
Rarefaction curves for each sample collected in the Baltic Sea Proper.

**Table S1.** Non-linear least squares analysis and statistics of observed colonization (*C*) and extinction (*E*) rates fitted with Hanski’s model (16) of *C*=c*P*(1-*P*) and *E*=e*P*(1-*P*). Asterisks denote significance levels (*P<0.05, **P<0.01, ***P<0.001) for NLS tests.

| Year | Month | Colonization ($C$) | | | Extinction ($E$) | | |
| --- | --- | --- | --- | --- | --- | --- | --- |
| | | Parameter Estimate | $p$ | Residual std. Error | Parameter Estimate | $p$ | Residual std. Error |
| 2010 | April | 0.23 | *** | 0.06 | -0.77 | *** | 0.05 |
|  | Early May | 0.39 | *** | 0.08 | -0.75 | *** | 0.04 |
|  | Late May | 0.31 | *** | 0.07 | -0.76 | *** | 0.04 |
|  | June | 0.44 | *** | 0.12 | -0.85 | *** | 0.06 |
|  | September | 0.28 | *** | 0.11 | -0.93 | *** | 0.08 |
| 2011 | April | 0.21 | *** | 0.06 | -0.86 | *** | 0.06 |
|  | May | 0.28 | *** | 0.11 | -1.06 | *** | 0.07 |
|  | June | 0.11 | *** | 0.07 | -1.05 | *** | 0.06 |
|  | July | 0.25 | *** | 0.09 | -1.00 | *** | 0.06 |
|  | August | 0.35 | *** | 0.09 | -0.74 | *** | 0.05 |
|  | September | 0.32 | *** | 0.09 | -0.82 | *** | 0.05 |

**Table S2.** Non-linear least squares analysis and statistics of observed extinction (*E*) rates fitted with Levin’s model (27) of *E*=e*P*. Asterisks denote significance levels (*P<0.05, **P<0.01, ***P<0.001) for NLS tests.

| Year | Month | Parameter Estimate | $p$ | Residual std. Error |
| --- | --- | --- | --- | --- |
| 2010 | April | -0.27 | *** | 0.09 |
|  | Early May | -0.75 | *** | 0.09 |
|  | Late May | -0.24 | *** | 0.09 |
|  | June | -0.29 | ** | 0.10 |
|  | September | -0.33 | ** | 0.12 |
| 2011 | April | -0.34 | ** | 0.12 |
|  | May | -0.46 | ** | 0.12 |
|  | June | -0.44 | ** | 0.12 |
|  | July | -0.47 | ** | 0.12 |
|  | August | -0.31 | ** | 0.09 |
|  | September | -0.31 | ** | 0.10 |

**Table S3.**
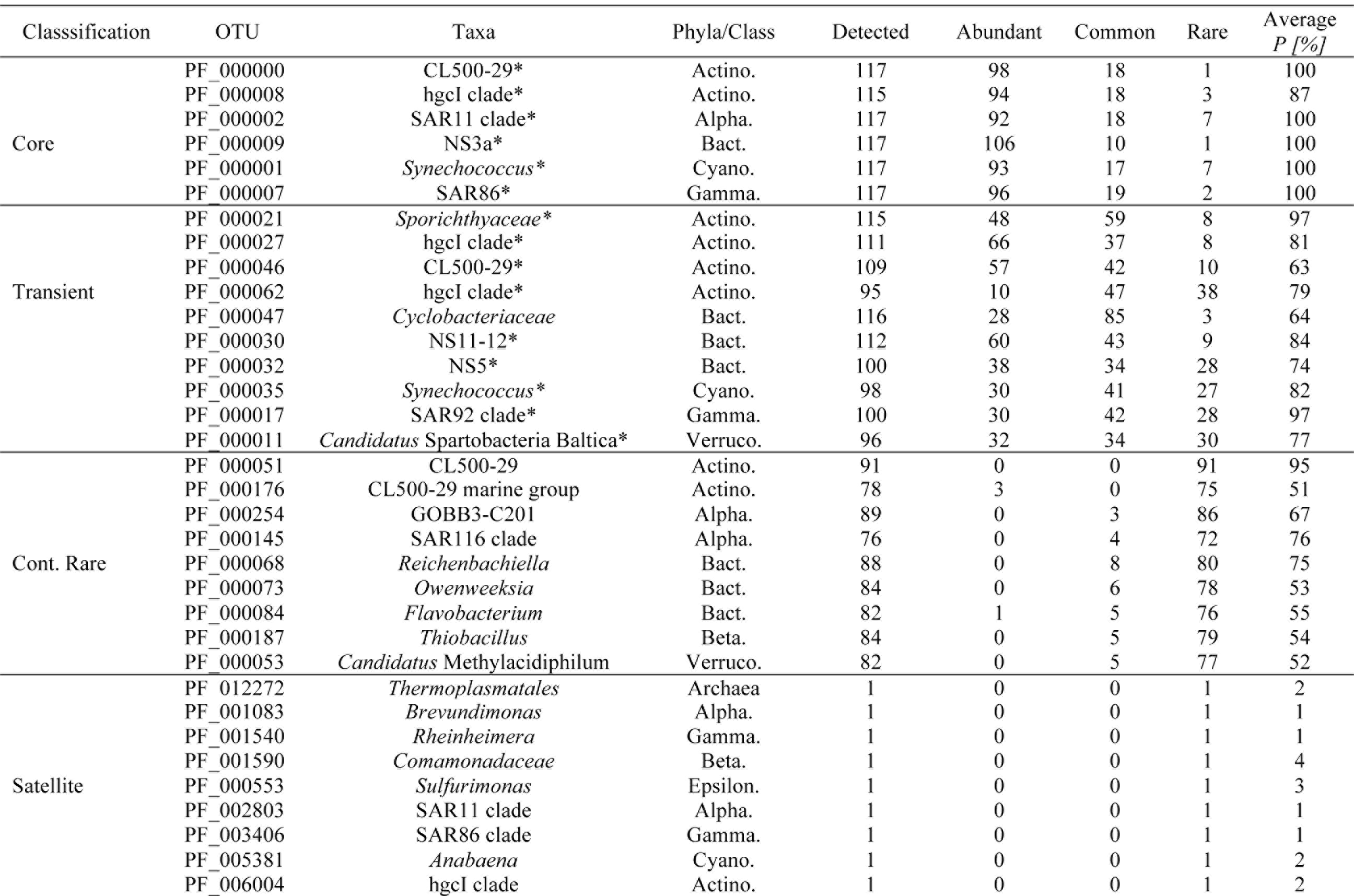
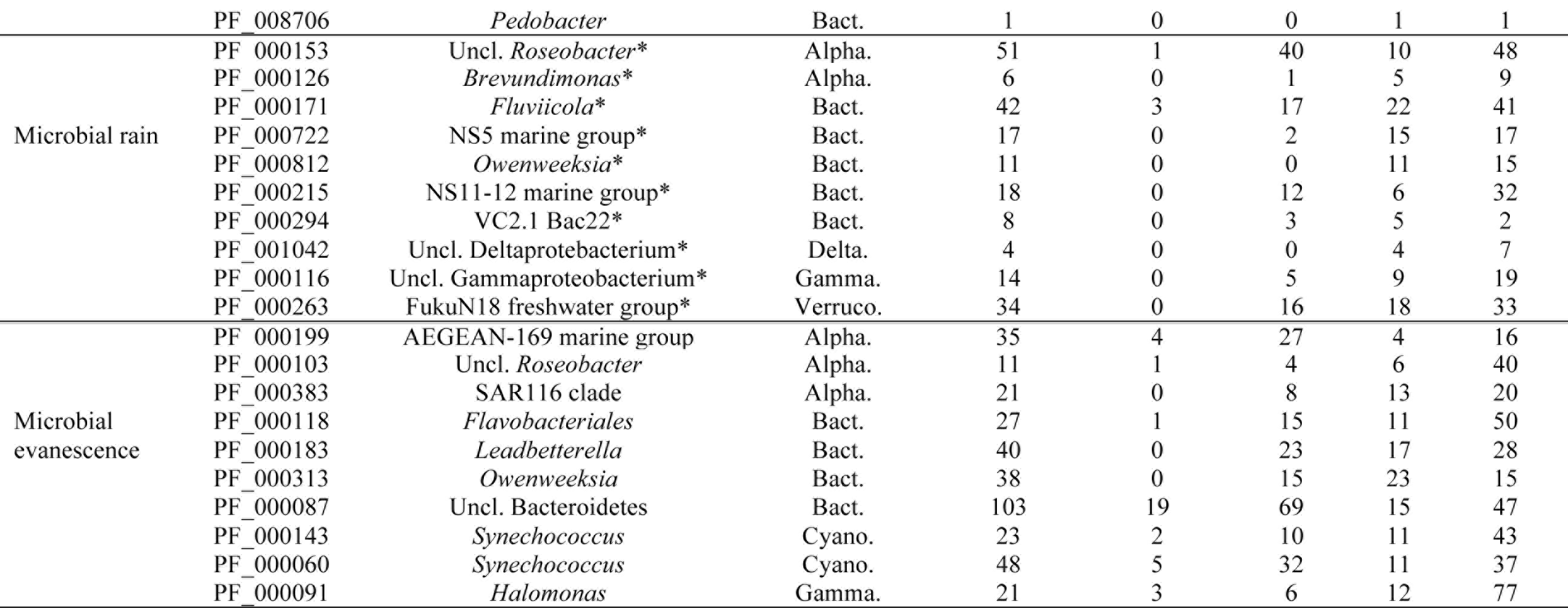
Classification of taxa into “core”, “transient”, “continuously rare”, “satellite”, “microbial rain” and “microbial evanescence” populations. Times detected of each individual OTU with times abundant (i.e. >1%), common (i.e. 0.1%-1%), rare (<0.1%), and average occupancy (*P*) in percentage. Populations with high average relative abundance > 1% and high occupancy with little variance in occupancy and relative abundance were classified as “core”. Populations with high average relative abundance and high variance in occupancy and relative abundance were classified as “transient”. Populations with low relative abundance but high occupancy and little variance in relative abundance and occupancy were classified as “continuously rare”. Populations with low average relative abundance and low occupancy were classified as “satellite”. “Microbial rain” and “Microbial evanescence” populations have large variance in occupancy, but have at one or more occasions gone from being undetected at all sites to being present at all sites, and vice versa, respectively. Asterisk (*) indicates if an OTU has been part of the rightmost bar in the bimodal analysis (Fig. 2) at least once.

